# WMP: A novel comprehensive wheat miRNA database, including related bioinformatics software

**DOI:** 10.1101/024893

**Authors:** Mohamed Amine Remita, Etienne Lord, Zahra Agharbaoui, Mickael Leclercq, Mohamed A. Badawi, Vladirmir Makarenkov, Fathey Sarhan, Abdoulaye Baniré Diallo

## Abstract

MicroRNAs (miRNAs) are emerging as important post transcriptional regulators that may regulate key plant genes responsible for agronomic traits such as grain yield and stress tolerance. Several studies identified species and clades specific miRNA families associated with plant stress regulated genes. Here, we propose a novel resource that provides data related to the expression of abiotic stress responsive miRNAs in wheat, one of the most important staple food crops. This database allows the query of small RNA libraries, including *in silico* predicted wheat miRNA sequences and the expression profiles of small RNAs identified from those libraries. Our database also provides a direct access to online miRNA prediction software tuned to *de novo* miRNA detection in wheat, in monocotyledon clades, as well as in other plant species. These data and software will facilitate multiple comparative analyses and reproducible studies on small RNAs and miRNA families in plants. Our web-portal is available at: http://wheat.bioinfo.uqam.ca.

## 1 Introduction

Next-generation sequencing offers interesting tools for generating and searching for microRNAs (miRNAs) that are expressed during growth and development. This approach provides several small RNA libraries and miRNA candidates from hexaploid wheat (*Triticum aestivum L.*) grown under different stress conditions such as heat, cold, and powdery mildew and *fusarium* infection. MiRBase, the current general database of miRNA sequences, is the main resource to access miRNAs data. Unfortunately, only 119 experimentally validated wheat miRNAs are available in the latest release of miRBase (Kozomara and Griffiths-Jones, 2011). Furthermore, the miRBase interface is not suitable for the analysis and assessment of miRNA candidates according to their original biological libraries and experimental conditions. To circumvent this limitation, we developed a novel comprehensive web-based server dedicated to wheat miRNAs, with emphasis on stress responsive miRNAs from different wheat genotypes and tissues. This database stores and displays differential gene expression data from several small RNA libraries. It also includes the data from 20 different miRNA studies (references are listed in the database interface). Moreover, the web portal yields and integrated views of the pre-miRNA hairpin structures and the related predicted targets. It also includes two miRNA predictors: *miRdup* (Leclercq *et al*., 2013) and *MIRcheck* (Jones-Rhoades *et al*., 2006) tuned for different evolutionary clades (wheat, cereals or plants).

## 2 Database content and statistics

The current database includes data of ten small RNA libraries produced from plants grown under different abiotic stress conditions and development stages. From these libraries, a set of 168,834 unique expressed small RNAs are illustrated as well as 267 evolutionary conserved miRNAs according to miRBase (Kozomara and Griffiths-Jones, 2011). The database also contains a broad description of 5,036 published wheat miRNAs. It also includes detailed presentation of 199 newly identified wheat miRNAs, 1,390 associated target genes, and 1.4 millions EST with 127,039 Uniref clusters, collected from seven wheat databases issued from Agharbaoui *et al*. (2015). Putative target genes were identified for each miRNA using the Tapir program (Bonnet *et al*., 2010). Gene ontology (GO) associations and enrichments were also performed for associated targets resulting of 2,561 biological process, 1,616 cellular components, and 1,386 molecular function associations. To enhance the visualization of miRNA structures, a high quality picture of each miRNA and each pre-miRNA hairpin were generated using the Varna package (Darty *et al*., 2009). Figure 1 highlights an example of graphical and statistical features of miRNA candidate in our database.

**Fig. 1.**
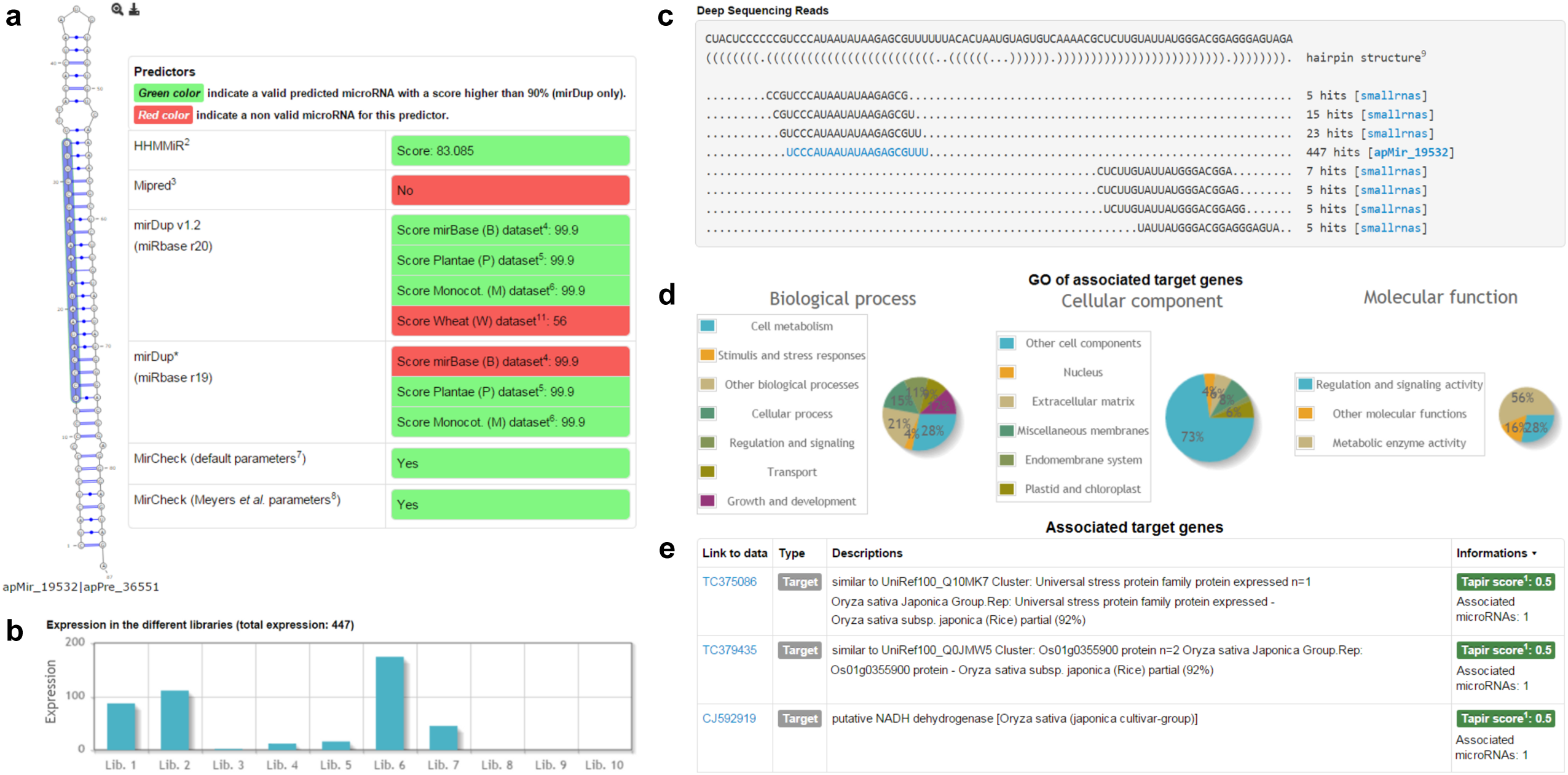
Overview of the main features of our miRNA database including (**a**) two-dimensional hairpin structures and the associated prediction scores for the miRNA apMir 19532, (**b**) expression view of each library, (**c**) view of small RNA expression and miRNA position against the computed pre-miRNA, (**d**) Gene Ontology annotations associated with a set of target genes and (**e**) description of the obtained target genes.

## 3 User interfaces

The database users have the following four main options: A) *Basic search*: the user can use keywords that correspond to one or multiple entries from the following categories: validated miRNAs, pre-miRNAs or the hairpin-folded structure motifs (dotbracket notation), associated target gene accession identifier, Uniref identifier or name, EST sequences or identifications, and GO descriptions or identifiers (e.g. GO:0008152). B) *Advanced search*: the user can search for differentially expressed miRNAs by choosing the growth conditions and selecting a list of miRNA patterns (upregulated, downregulated, not differentially expressed, or not related to any of the conditions). The metrics for statistical significance (*p-values*) and the miRNA expression fold-change between libraries could also be set to filter candidates. C) *Data* menu: provides a direct access to predicted miRNAs, conserved miRNAs, the associated target genes and ESTs. This menu also gives access to *Libraries and conditions* option. This option allows the profiling all miRNAs expressed in any given libraries or under given stress conditions. D) *Tools* menu: provides five important applications for studying small RNAs libraries and miRNAs. It includes *MIRcheck v1.0* (Jones-Rhoades and Bartel, 2004), *miRdup v1.2* (Leclercq *et al*., 2013), *Small RNA finder*, *Library comparison* as well as a *Blast* interface (Altschul *et al*., 1990; Camacho *et al*., 2009). *MIRcheck* and *miRdup* are miRNA prediction tools. For both predictors, the users can submit their candidate miRNAs, miRNA precursors, or hairpin secondary structures for analysis. The *MIRcheck* tool allows two modes of computation (default parameters used by Jones-Rhoades and Bartel (2004) and the universal plant miRNA criteria given by Meyers *et al*. (2008)). The *miRdup* predictor (based on a machine learning approach) allows a selection among four evolutionary clade-training sets. These sets are extracted from experimentally validated miRNA sequences found in miRBase (all miRNAs of miRBase, viridiplantae (plant), monocotyledons and wheat (*Triticum aestivum L.*)). *Small RNA finder* option permits users to search multiple sequences in a single query against putative miRNA and small RNA database. *The library comparison* option allows extracting differential expressed small RNAs between two libraries. In the *Blast* option, users can identify the location of their query sequences (nucleotide and amino acid sequences) against the wheat whole sequenced genome using the *Blast* tool. Further details on our database and a demonstration of some of its features are provided in the following section.

## 4 Study case: searching for miRNAs regulating glutathione S-transferases

The usefulness of our miRNA database is shown with an example of the search of miRNAs that regulate glutathione S-transferase (GST) enzyme family in wheat. GSTs are multifunctional proteins known for their important roles in both normal cellular metabolisms and the detoxification of a wide variety of products under stress conditions (Marrs, 1996; Jain *et al*., 2010). GSTs have been shown to be miRNA targets in various other plant species exposed to stresses, such as osa-miR1848 in rice (Li *et al*., 2010; Shaik and Ramakrishna, 2012), rsa-miR156 in radish (Xu *et al*., 2013), mir168 homolog in sweet potato (Dehury *et al*., 2013) and ssp-mir169 in sugarcane (Gentile *et al*., 2013; Menossi *et al*., 2015). Using the basic keyword search option, we first looked for the complete name of the GST enzyme: glutathione S-transferase. We found 31 target gene sequences, one microRNA gene and 30 gene ontology annotations. To ensure that we didn’t miss any other related data in the database we searched for alternative keywords related to the enzyme such as glutathione transferase and GST. Clicking on the target sequence identifier CJ893705 (with a good Tapir score of one), allows the user to access further related information such as the alignment with the miRNA apMir 11506, the Uniref clusters where the target belongs and the associated GO annotations. Information related to the found miRNA such as its sequence, length and expression details, can be easily obtained by clicking on its identifier link. In the *in silico* evidence section of the miRNA page, one could find that it is embedded in two precursors (apPre 12405 and apPre 8495) and it is predicted as valid miRNA with two different predictors *miRdup* ((Leclercq *et al*., 2013) and *MIRcheck* (Jones-Rhoades and Bartel, 2004)). A deep sequencing reads view of precursors and their mapping small RNAs is appended to the evidence section. Interestingly, we noticed that apMir 11506 is regulated by cold and salinity. Furthermore, in miRBase section, we found that it has a 20-nucleotides homolog miRNA in Arabidopsis thaliana (ath-miR8175). These results show that in total, 5 miRNAs are associated with glutathione S-transferase: apMir 54471, apMir 15117, apMir 18589, apMir 11506 and apMir 19203. The first four miRNAs target the GST enzyme. apMir 18589 and apMir 19203 are generated by ESTs associated with GST Uniref. Analyzing the expression behavior in the different conditions reveals that miRNAs targeting the GST were regulated in aluminum (Al), salt and cold stresses from sensitive and tolerant wheat genotypes.

## 5 Conclusion

The WMP database presents a novel resource for the analysis of miRNAs in cereals. This first version offers an access to miRNAs and small RNAs in the context of their association with different growth and development stages under stress conditions in wheat. It constitutes a unique entry for wheat expressed small RNAs and miRNAs for several studies. Furthermore, it provides a direct access to useful miRNA predictors for wheat species, as well as for other cereals and plants. In the future, we plan to update this database regularly by adding new libraries prepared by our team for wheat grown under different stress conditions and newly published small RNAs in wheat and other cereals. We will provides confidence metrics over all the published miRNAs in wheat. We will also includes a evolutionary toolkits to study miRNAs within all cereals. Further development will include option to integrate complex miRNA analysis pipelines using the Armadillo workflow platform (Lord *et al*., 2012).

## 6 acknowledgements

Acknowledgements This work was supported by the Natural Sciences and Engineering Research Council of Canada (NSERC) to FS and ABD and the Fond de Recherche du Qubec-Nature etTechnologie (FRQNT) to ABD. MAR is a NSERC and FRQNT fellow.

